# Photosynthesis is heavily chlororespiration-sensitive under fluctuating light conditions

**DOI:** 10.1101/151530

**Authors:** Wojciech J. Nawrocki, Felix Buchert, Pierre Joliot, Fabrice Rappaport, Benjamin Bailleul, Francis-André Wollman

## Abstract

Photosynthesis needs to adjust to dynamically changing light intensities in order to maximize its efficiency, notably by the employment of alternative electron pathways. One of them is chlororespiration - initially described in *Chlamydomonas reinhardtii*. This electron transfer pathway, found in all photosynthetic lineages, consists of a reduction of plastoquinone (PQ) through an NAD(P)H:PQ oxidoreductase and quinol (PQH_2_) oxidation by Plastid Terminal Oxidase, PTOX. Hence, chlororespiration constitutes an electron pathway potentially antagonistic to the linear photosynthetic electron flow from H_2_O to CO_2_. However, the limited flow chlororespiratory enzymes can sustain suggests that their relative contribution, at least in the light and in steady-state conditions, is insubstantial. Here, we focused on the involvement of PTOX in *Chlamydomonas reinhardtii* during transitions from dark to light and vice versa. We show that the kinetics of redox relaxation of the chloroplast in the dark was greatly affected when PTOX2, the major plastoquinol oxidase in *Chlamydomonas*, is lacking. Importantly, we show that this has a direct physiological relevance, as the growth of a PTOX2-lacking mutant is markedly slower in intermittent light. The latter can be rationalized in terms of a decreased flux sustained by photosystem II due to a redox limitation at the acceptor side of the PSI during the illumination periods. We finally show that the long-term regulation of cyclic electron flow around PSI is strongly affected in the PTOX2 mutant, substantiating an important role of chlororespiration in the maintenance of chloroplast redox balance.

## Introduction

Plants need to cope with large variations in light irradiance in order to maintain efficient photosynthesis. There are multiple regulatory processes in the chloroplast that contribute to optimize carbon fixation via linear electron flow (LEF) from photosystem II (PSII) to the Calvin Benson Basham (CBB) cycle. They also provide protection against over-reduction of the electron transport chain, which otherwise would lead to the production of deleterious reactive oxygen species (1-3). These include two major processes: (i) a downregulation of PSII photochemical activity either by quenching of its excitonic energy (energy-dependent non-photochemical quenching, called q_E_)(4) or by decreasing its absorption cross-section (state transitions, ST/q_T_)(5, 6); and (ii), a rerouting of part of the electron flow from PSII to alternative sinks, mostly oxygen (water-to-water cycles such as Mehler reaction photorespiration, or flavo-diiron protein activity)(7). Cyclic electron flow (CEF) around PSI, which reinjects electrons from reduced ferredoxin back to plastoquinone (PQ) and thereby increases the proton gradient across the thylakoid membrane, contributes both to the downregulation of the PSII and PSI activity and to the optimization of the LEF by providing extra ATP to the CBB cycle (8, 9).

The aforementioned mechanisms regulate electron flow in the light, yet there exist other regulatory processes, which are light-independent but share part of the electron transport chain. One of these is chlororespiration, defined as a light-independent oxygen consumption in chloroplasts (10). It is now well established that chlororespiration is conserved in the photosynthetic lineage and involves two types of enzymatic activities that act in series and share the substrate, the plastoquinone/plastoquinol pool. An NAD(P)H:PQ oxidoreductase (NDA2 in algae, NDH in higher plants) reduces the plastoquinones using stromal reductants as an electron donor (11), and a PQH_2_:O_2_ oxidoreductase termed PTOX (for Plastid Terminal OXidase) which oxidises plastoquinols (12) and produces water. Altogether, the in-series activity of NDA2/NDH and PTOX transfers electrons from the stromal reductants to O_2_, and their differential amounts and kinetic constants regulate the redox state of the PQ pool in the dark (13). In higher plants, PTOX is also involved in carotenoid biosynthesis during chloroplast development, but its presence in adult organisms suggests that it continues to play a physiological role (14, 15). This role, however, has remained elusive up to now.

Chlororespiration is thus intertwined with the photosynthetic electron transfer chain by sharing PQ and some stromal electron carriers. As an antagonist of the light reactions of the photosynthesis, it consumes the primary products O_2_ and NADPH (12). We previously have shown that a mutant devoid of the major chlororespiratory PQ oxidase in the green alga Chlamydomonas (PTOX2) exhibits no difference in fitness when grown in the dark in mixotrophic conditions even though its PQ pool remains fully reduced (13). The maximal rates of the chlororespiratory enzymes are – depending on the species – one or two orders of magnitude lower per chain unit than the rate of the slowest step of electron transfer, i.e., oxidation of a PQH_2_ at the Q_O_ site of cytochrome *b*_6_*f* (cyt. *b*_6_*f*) (13, 16). These low rates suggest that chlororespiratory fluxes cannot compete with the main PQ reductase (PSII) or oxidase (cyt. *b*_6_*f*) of the photosynthetic chain in the light. Indeed, we show here that PTOX2 plays no significant role during steady-state illumination in Chlamydomonas. In marked contrast, we demonstrate that its activity becomes critical under fluctuating light conditions. We provide evidence that the chlororespiratory activity not only oxidises the PQ pool but also extensively oxidises stromal reductants in darkness, thereby modifying the efficiency of photosynthesis at the onset of light, while modifying the balance of the electron flows towards CEF or LEF.

## Results

### PTOX2 cannot compete with cyt. *b*_6_*f* - PSI in continuous light

Even though the low rate of PTOX is poorly suited for its role as an efficient electron sink (13), several studies suggested it had a photoprotective role when the light absorption by PSII exceeds the capacity of the Calvin-Benson-Bassham (CBB) cycle to reduce carbon (14, 17, 18). We first tested the significance of PTOX as a pathway rerouting electrons from PSII by monitoring the electron transfer rate (ETR) in the WT and ΔPTOX2 at different light intensities. Indeed, if PTOX accepts a significant proportion of the electron flux from PSII, then the ETR should be higher (by that amount) in the WT than in the mutant. As shown in the Fig. 1, the presence of the major PQH_2_:O_2_ oxidoreductase PTOX2 in Chlamydomonas does not significantly modify the ETR in the steady-state even at low light intensities of about 10 μE.m^-2^.s-^1^ (inserts in Fig 1) in both auto- and mixotrophic conditions. This indicates that PTOX activity is not relevant under continuous illumination and calls into question the hypothesis about its photoprotective role (12).

**Figure 1.**
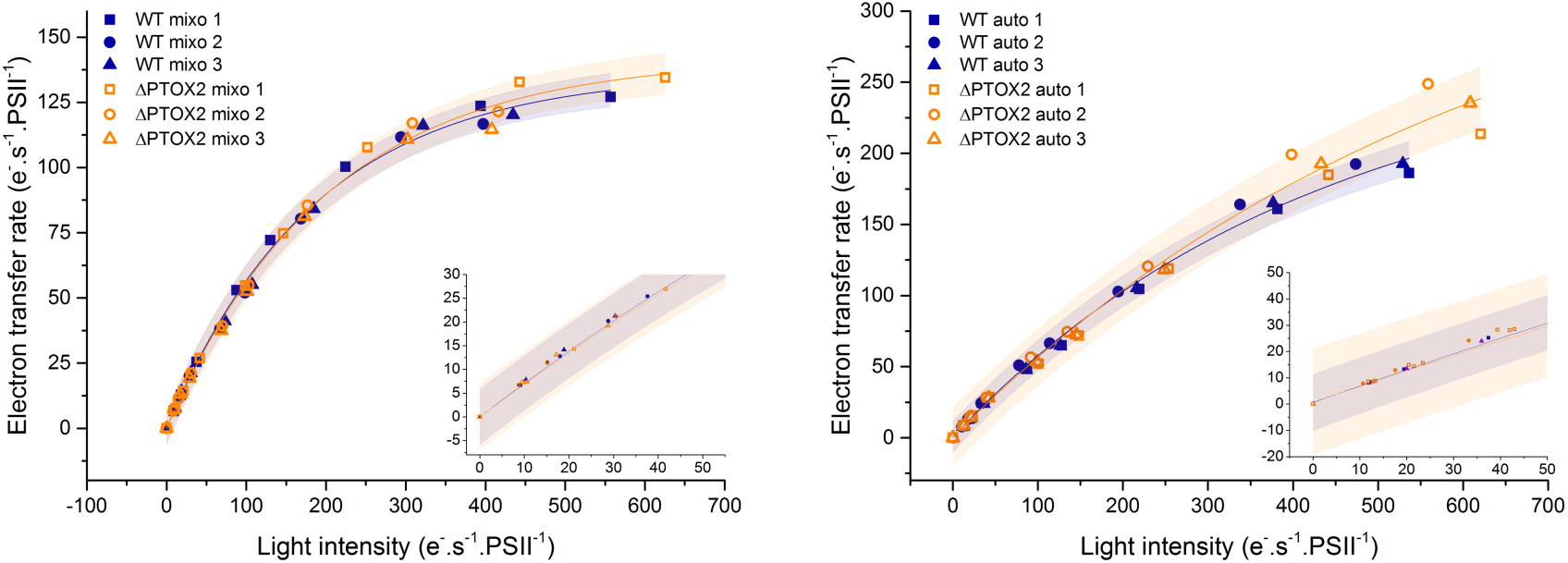
Electron transfer rates in WT and ΔPTOX2 grown in autotrophic (auto) and mixotrophic (mixo) conditions over a range of light intensities. ETR is calculated from the ΦPSII parameter after 30 s adaptation to each light intensity. See Materials and Methods section for details. *n* = 3 ± 95% confidence band of the exponential fit. Insets show the values for low light intensities.

### The ΔPTOX2 mutant exhibits severe growth and photosynthesis phenotypes in fluctuating light

Despite the absence of growth phenotype of the PTOX mutant when kept in darkness or under continuous illumination, we observed that its growth was severely altered in conditions where those two situations alternate (fluctuating light). As shown in Fig 2, the growth rate of the ΔPTOX2 strain severely decreased with regards to that of the WT in light conditions corresponding to cycles of 1 minute darkness followed by 1 minute illumination. This proved to be the case in both mixo- and autotrophic conditions, although the differences were less pronounced in the latter condition.

**Figure 2.**
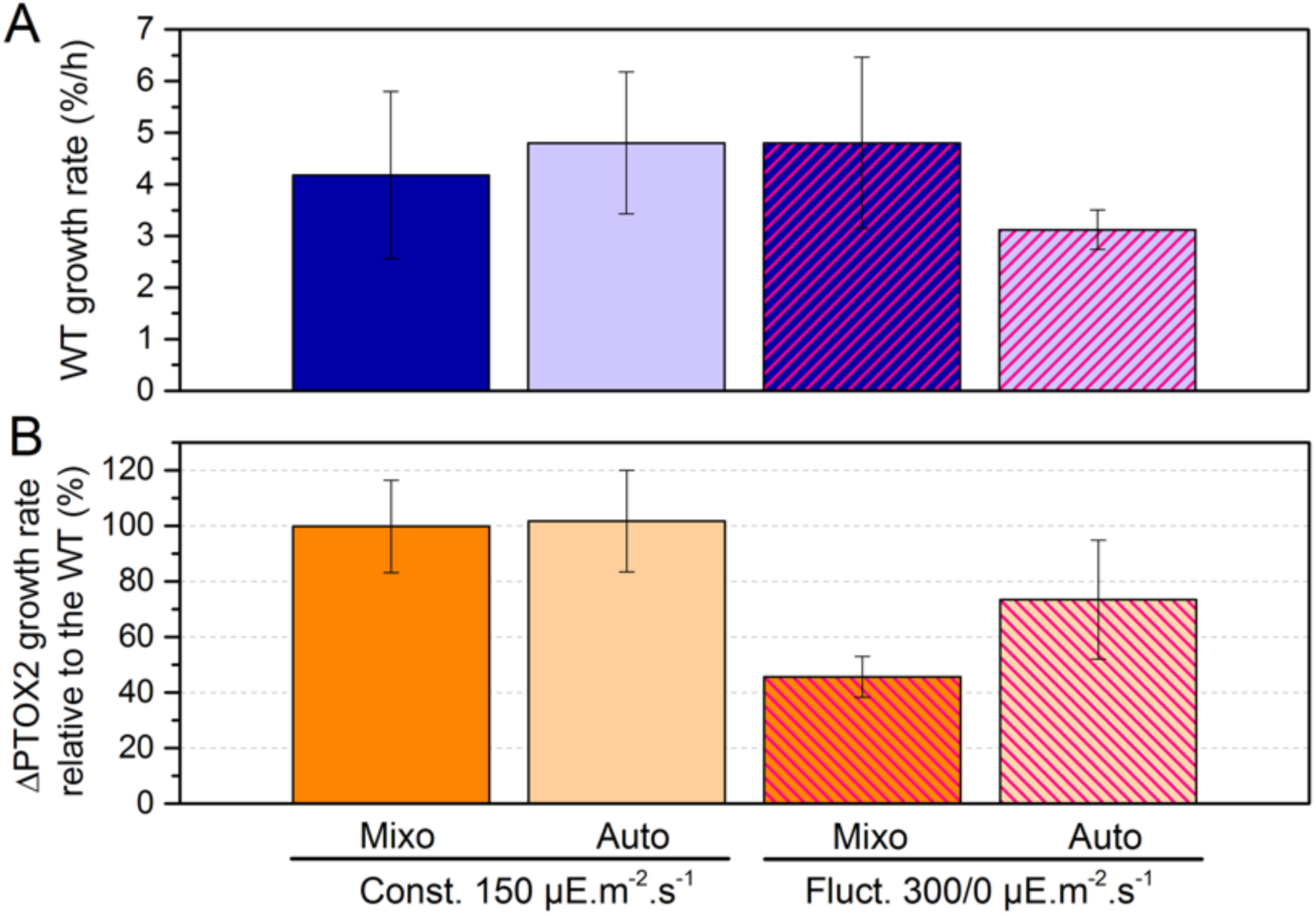
Growth rates of the WT (A) and relative growth rate of the ΔPTOX2 (B) in constant and fluctuating light conditions. (*n* = 3 – 5 ± S.E.)

These fluctuating light conditions were accompanied by an increase in the amount of PTOX2 in wild type cells, with the strongest change observed in mixotrophic growth after 5 h exposure to fluctuations (Fig. S6). A specific contribution to the growth behaviour in fluctuating light could arise from a labile and light-dependent binding of PTOX to the thylakoid membranes. It has been recently reported that PTOX would detach from higher plant thylakoids in the dark, supposedly due to a lower stromal pH. It would thus become stroma-soluble and unable to oxidise quinols (20). This behaviour could have an impact on the bioenergetics of the chloroplast in fluctuating light. Therefore, we examined whether changes in thylakoid binding of PTOX could develop between dark and light exposure. As shown in Figure S1, we did not observe any change in PTOX binding to the membranes between light and darkness. In the two cases, PTOX was found virtually exclusively in the membrane fraction.

We then measured the photochemical yield of PSII (ΦPSII) in cells grown in fluctuating light conditions to assess whether a change in photosynthetic electron fluxes could account for these differences in growth rates (19). As shown in Fig. 3, the decrease of ΦPSII in the ΔPTOX2 paralleled the decrease in growth rate of the mutant. These differences largely relaxed rapidly (in only 15 minutes) when we used a continuous illumination period after the fluctuating light regime (Fig. 3C). The latter observation shows that the growth phenotype is indeed due to a hampered capacity to respond to fluctuating light and not to changes in the architecture/composition of the photosynthetic apparatus in the course of the growth experiment.

**Figure 3.**
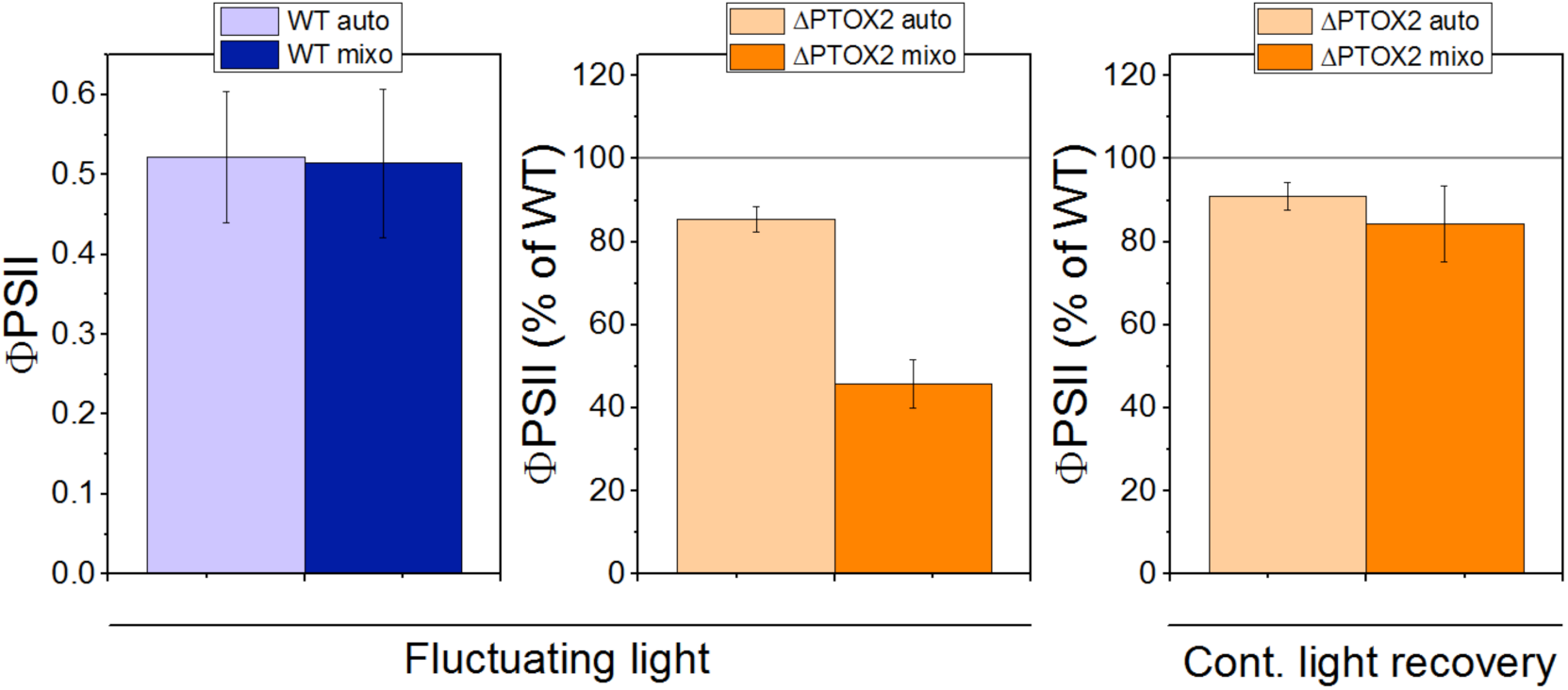
ΦPSII of WT and ΦPSII of ΔPTOX2 grown in fluctuating light, relative to the WT. (*n* = 3 – 5 ± S.E.).

We then investigated the influence of PTOX during both dark- and light periods on the whole photosynthetic process. In order to follow the short-term response of photosynthesis to the fluctuating light conditions, we placed the two strains in the absorption difference spectrophotometer under the same light fluctuations that they experienced during growth (see Materials and Methods). Given the role of the redox state of the PQ pool in the regulation of state transitions (13), WT and mutant cells were first adapted to low light intensity, setting both strains in state I prior to the fluctuating light treatment (fig. S2), a feature not achievable in darkness (20). The fluctuating light treatment in the spectrophotometer resulted in a similar phenotype as observed during growth conditions, i.e., a ΦPSII drop to a similar extent (fig. S3A), being reversible under continuous illumination.

We simultaneously recorded the evolution of the F_M_ and F_V_/F_M_ parameters during a 30 min exposure to fluctuating light (fig. S3B, C). No changes were observed between the strains, suggesting that neither q_E_ nor q_T_ differed between the WT and the ΔPTOX2 (see also Fig. S2). We then checked whether the ΦPSII phenotype of the mutant depended on the light intensity and periodicity of the fluctuating light regime. Using a lower (125 μE.m^-2^.s^-1^) or higher (1500 μE.m^-2^.s^-1^) light intensity (fig. S3D, E), and lower (20 s dark / 20 s light) or higher (180 s dark/ 180 s light) frequency (fig. S3F, G), we observed a similar drop in ΦPSII in the mutant compared to the WT. Last, we investigated the evolution of ΦPSII and F_V_/F_M_ parameters in a 1s dark / 1s light fluctuating light regime (fig. S4). Here, the differences were no longer observed, indicating that slow processes were responsible for the phenotype, which was in agreement with the low rate at which PTOX operates. All the experiments above were performed using cells grown in mixotrophic conditions. However, similar observations were recorded using cells grown in autotrophic conditions, although to a slightly lesser extent (fig. S5).

### Chlororespiration mediates the redox state of both the PQ pool and the PSI acceptors during dark periods

To understand how processes occurring in the dark can have consequences on the photosynthetic electron transfer during the light period, we investigated the redox relaxation of the photosynthetic electron transfer chain during the dark periods. We first explored the rate of Q_A_ re-oxidation during the first and last dark period of the 30 min treatment. We applied the Stern-Volmer relationship to quantify changes in the concentration of Q_A_^-^ from the fluorescence data (see supplementary materials for the equations) after applying a saturating pulse to reduce 100% of Q_A_ (21). As shown in the figure 4A, the kinetics of Q_A_ re-oxidation, reflecting the redox state of the PQ pool that is in equilibrium with Q_A_ in the dark, were very different in the two strains. In the absence of PTOX2, the oxidation of the PQ pool was less efficient and 40% of PSII centres remained in a partially reduced state at the end of the 1 minute dark period. This is in marked contrast to the situation in the wild type, in which the quinones were completely re-oxidized after 1 minute in the dark. The biphasic Q_A_ oxidation kinetics observed in the two strains were consistent with the fast phase reflecting the cyt. *b*_6_*f*-mediated re-oxidation of PQH_2_ by oxidized plastocyanin, and the slower phase corresponding to PTOX-mediated re-oxidation of PQH_2_.

It is noteworthy that Q_A_ re-oxidation was slower after the 30-min. light/dark treatment in both strains, suggesting an accumulation of reductants under such regime that would translate in a higher level of reduction of the PQ pool and Q_A_. In fluctuating light, a hypothetical increase in the concentration of the NDA2 substrate, NAD(P)H, would be consistent with these observations. Accordingly, such an increase would be even more pronounced in the ΔPTOX2 strain due to decreased chlororespiration in the presence of the remaining minor PQH_2_:O_2_ oxidoreductase, PTOX1. Consistent with this view, Q_A_ remained still more reduced in the mutant (Fig. 4B).

**Figure 4.**
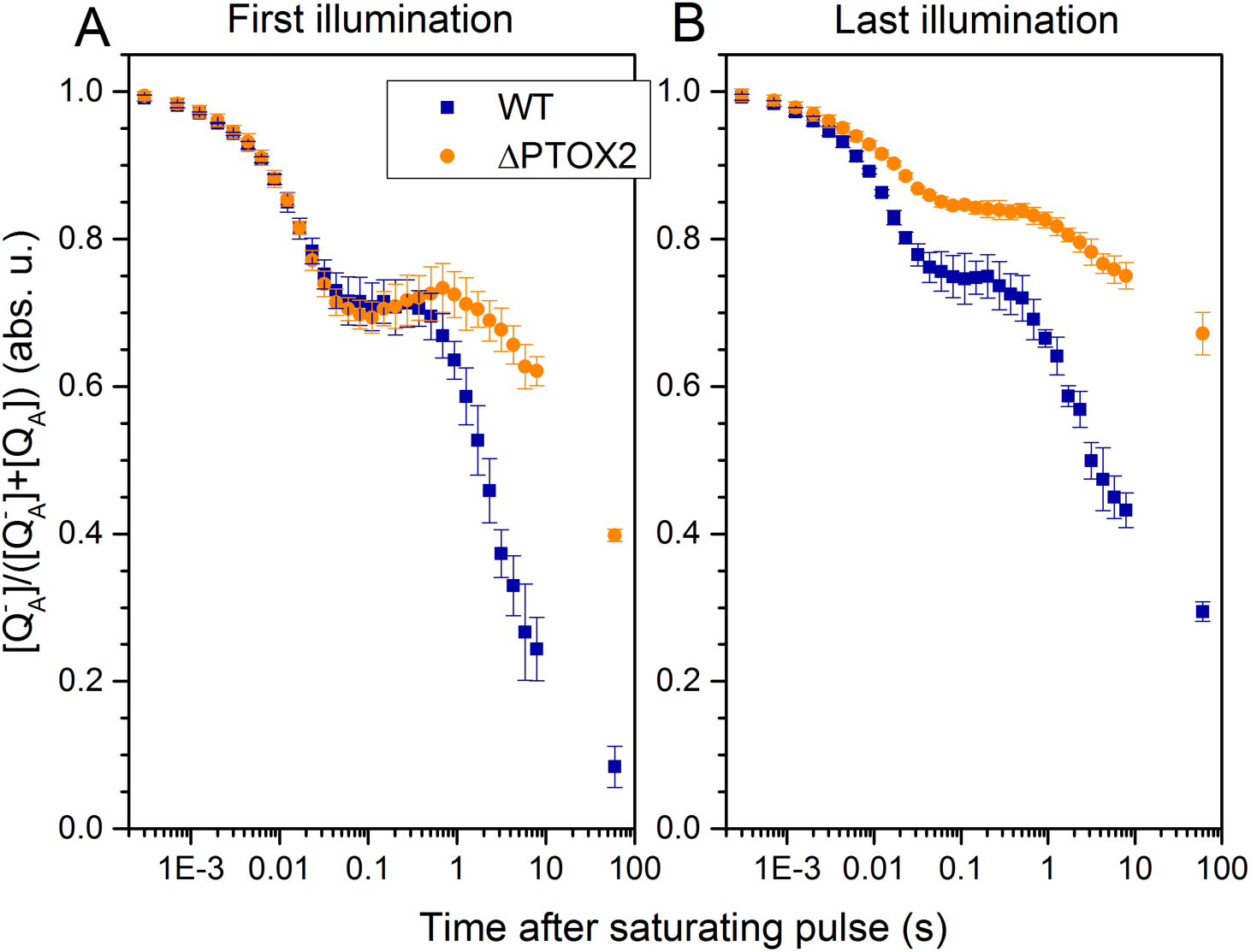
The kinetics of Q_A_^-^ concentration changes in darkness after a saturating pulse in fluctuating light as determined from the fluorescence decay kinetics. The maximal quantity of quencher (corresponding to the [Q_A_^-^] value of 0) reflected the low-light adapted condition, followed by a brief dark-adaptation. A, after the first illumination period; B, after 30 minutes of fluctuating light treatment. *n* = 3 ± SD

To further test this hypothesis, we measured the acceptor side limitation of PSI at the end of the dark period throughout the 30-min. treatment using the Klughammer and Schreiber protocol (22). The results are shown in the figure 5. The amount of photo-oxidisable P700 decreased in both strains, indicating an increase in PSI acceptor side limitation (accumulation of stromal reductants) with increasing exposure to fluctuating light treatment. This accumulation was stronger in the ΔPTOX2, as indicated by the steeper decrease of the photo-oxidisable P700. Therefore, we concluded that chlororespiration is necessary in darkness to recycle the stromal reductants produced during the light period.

**Figure 5.**
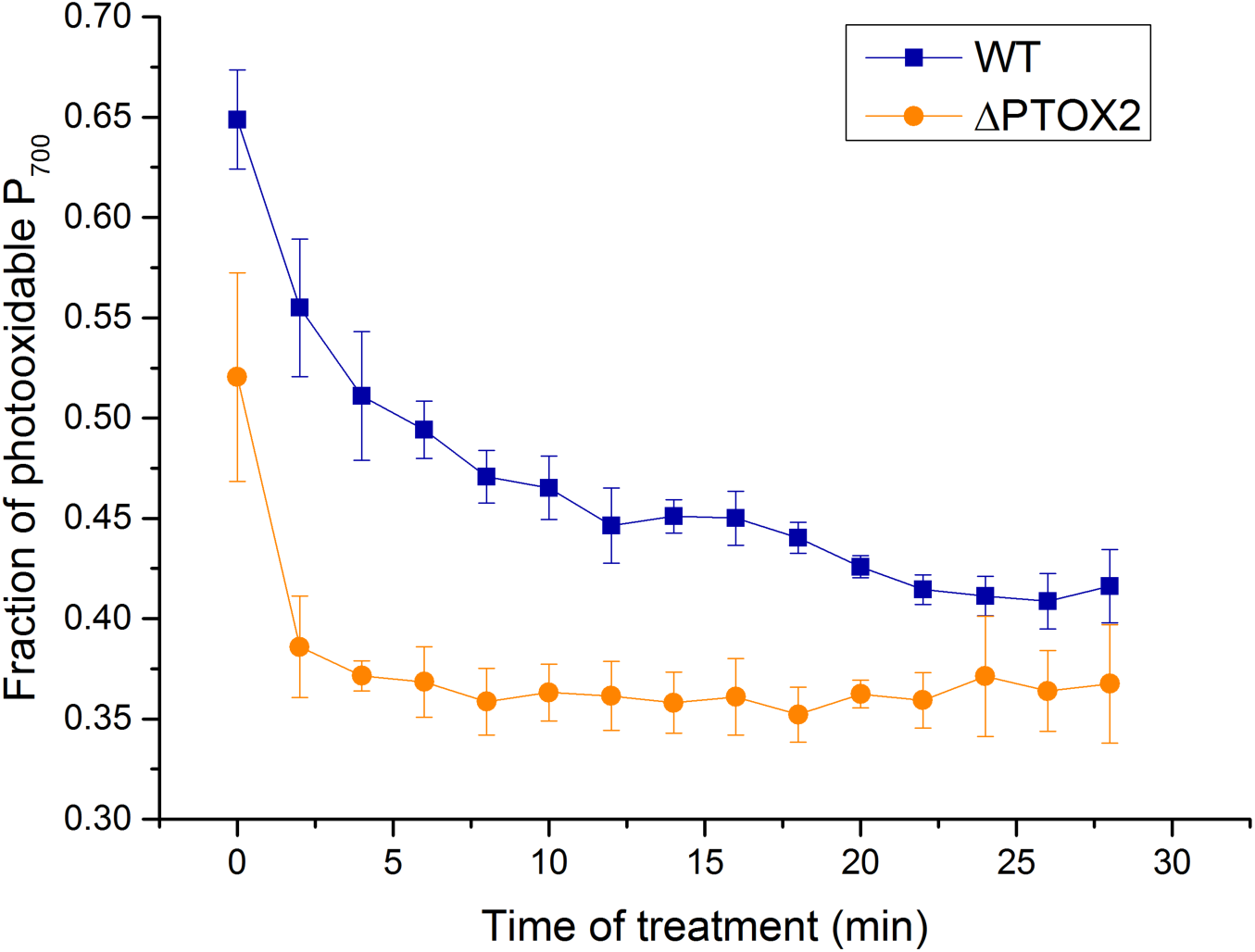
Effect of the fluctuating light treatment on the quantity of photo-oxidisable P_700._ The maximal quantity of P_700_^+^ probed during a 20ms saturating pulse at the onset of each light period was normalised to a DCMU-treated reference to yield a fraction of total P_700_. *n* = 3 ± SD

### Cyclic electron flow around PSI is upregulated in the absence of chlororespiration

We also followed the evolution of ΦPSII and ΦPSI at the end of each light period during the 30 min exposure to fluctuating light. As shown in figure 6, ΦPSII remained unchanged in the wild type while it markedly dropped in the ΔPTOX2 strain. However, ΦPSI increased in both strains during the same treatment. It reached the same value as that ΦPSII in the wild type, whereas it increased further in the PTOX2 mutant, reaching a value about two times larger than that of ΦPSII at the end of the treatment. We interpret these unbalanced photochemical yields in the mutant as an upregulation of cyclic electron flow (23), which indeed should increase PSI activity relative to PSII. Strictly speaking, ratios of PSII_to_PSI photochemical yields can only be interpreted as a ratio of PSII-to-PSI activities if the antenna sizes of PSII and PSI behave the same in the WT and the mutant. Since the decrease of F_M_ (reflecting both q_T_ and q_E_) and the amplitude of the state transitions (q_T_) were similar in the two strains, we conclude that the higher ratio of ΦPSII/ΦPSI reflects a higher CEF in the mutant (Figs S2, S3; see also supplementary discussion).

**Figure 6.**
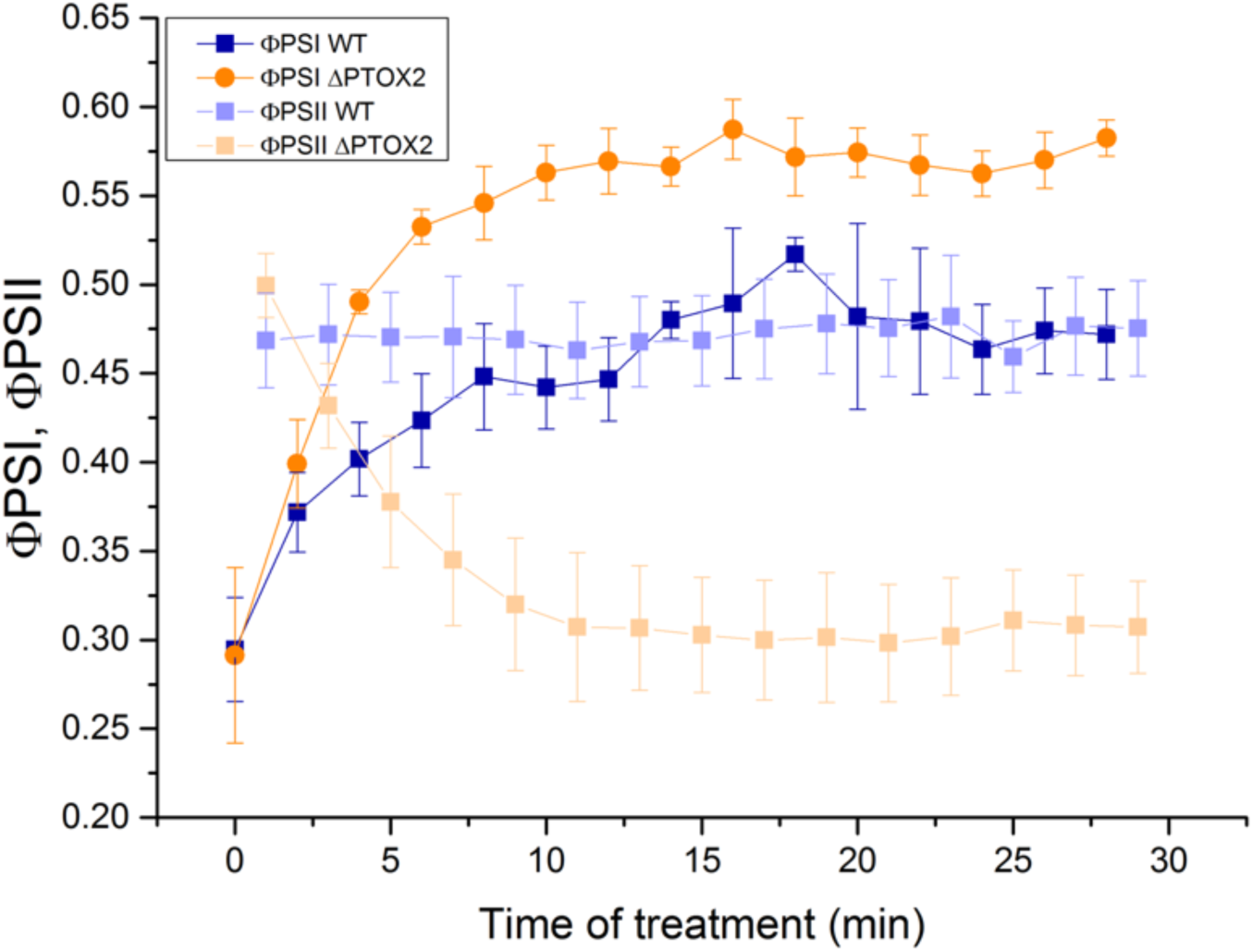
Evolution of the photochemical yields of PSII and PSI during the fluctuating light treatment. *n* = 4 ± SD

We then measured the total quantity of photo-oxidisable PSI at the end of each light period and the fraction of oxidised PSI accumulating during this light period to establish whether the over-reduction of the PSI acceptor side observed in Fig. 5 persisted throughout the light period. In contrast to the wild type, ΔPTOX2 accumulated a significant proportion of P700^+^ in the light together with a higher amount of photo-oxidisable P_700_ (Fig. 7). Thus, in the absence of PTOX2, the donor side limitation was higher while the acceptor side limitation was largely decreased. These observations suggest that the high CEF efficiency in absence of PTOX is not the mere result of a higher reduction of the pool of stromal electron carriers in the light, but a regulatory response at the level of thylakoid membranes, of which the molecular basis remains to be elucidated.

**Figure 7.**
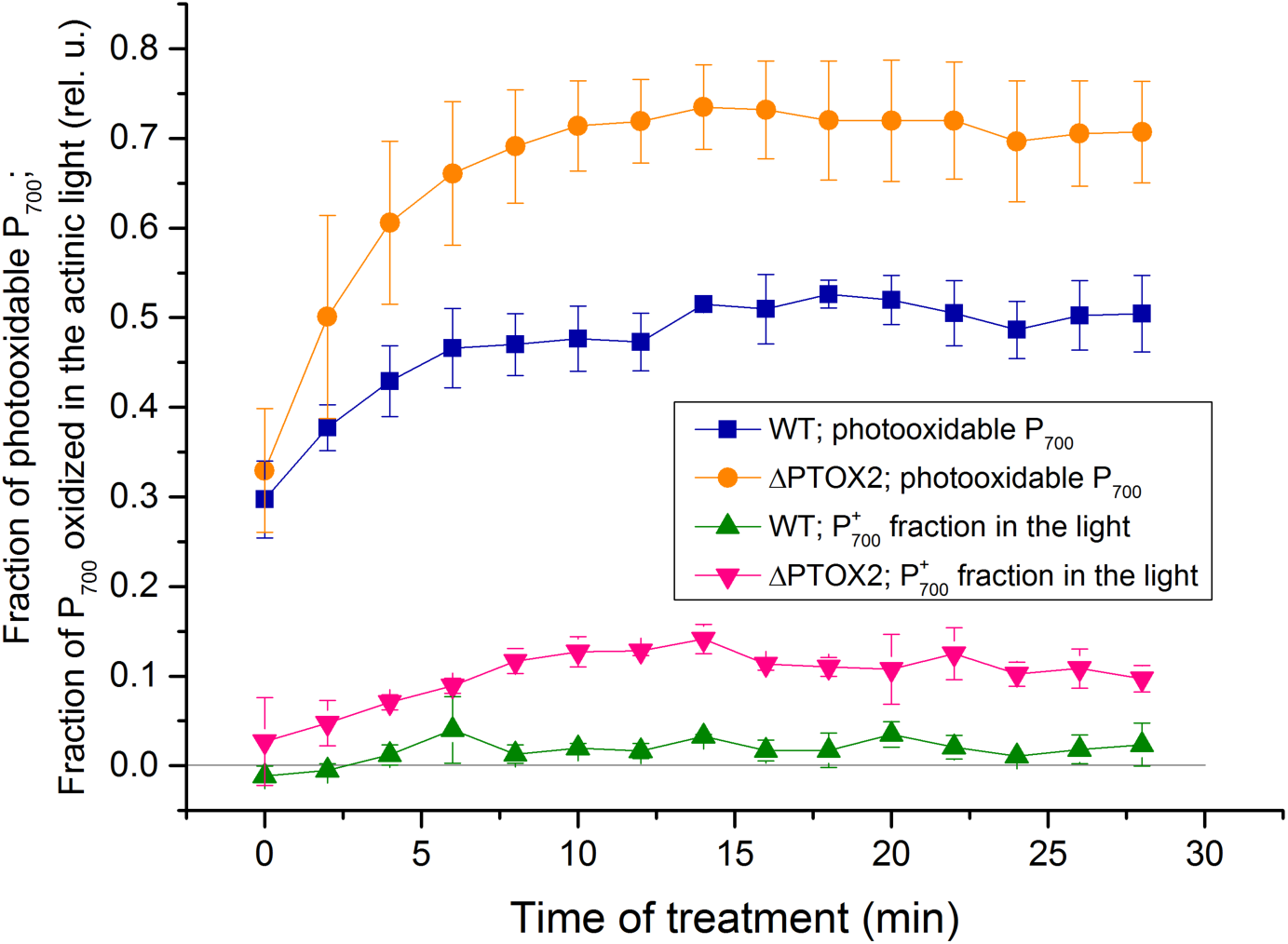
PSI redox state parameters during the fluctuating light treatment. Absorption at 705 nm was monitored during the light-to-dark transitions in the WT and in the PTOX2 mutant. Fraction of photo-oxidisable P700 was determined using a short saturating pulse at the end of the light period; P_700_^+^ accumulation in the light corresponds to the difference between absorption before the pulse and in darkness. Both quantities are shown as a fraction of total P_700_ that were determined in the DCMU-treated sample in the light with a saturating pulse superposed. *n* = 3 ± SD

## Discussion

Although discovered decades ago, chlororespiration had never been properly understood, and its role remained unknown. Little or no phenotypes of chlororespiration mutants hampered unravelling the role of this potentially futile process. However, to date the investigations to elucidate the role of chlororespiration were performed exclusively in constant light or in darkness. We show that the ΔPTOX2 mutant exhibits decreased growth rates with regards to the WT in fluctuating light, but neither in continuous light nor in darkness. In that respect, the PTOX enzyme resembles a number of other auxiliary proteins in photosynthesis whose physiological role becomes prominent under fluctuating light conditions but not in the standard laboratory conditions, such as constant light (24). In particular, Flavodiiron proteins seem to perform a similar function in cyanobacteria and Chlamydomonas where they serve as a typical overflow device – oxidising part of NAD(P)H when a saturating light is superposed on growth light (25, 26). Similarly, the regulation of electron flux mediated by state transitions is important for growth in fluctuating light or over-reducing conditions (27, 28). In cyanobacteria, multiple mutants of terminal oxidases exhibit a growth phenotype when exposed to fluctuating light at high light intensity (29), but the photosynthetic parameters that may be affected were not measured in these cultures. Similarly, photosynthesis in fluctuating light conditions in the CEF mutants was impaired (30) – the authors show that PSI is the target of photoinhibition in such case. Although we cannot rule out photo-oxidative modifications, in our hands the total amount of PSI did not decrease, despite PSI acceptor side over-reduction upon illumination which is usually connected to PSI photoinhibition.

We have shown that the growth differences in fluctuating light correlate with changes in the ΦPSII and F_V_/F_M_ chlorophyll fluorescence parameters. Their interpretation is that the PSII-driven electron transfer is strongly decreased in the ΔPTOX2, which in turn impedes growth of the algae. Importantly, we also show that chlororespiration strongly affects not only the PSI donor side, but also its acceptors. Therefore, it regulates the entire ambient redox potential of the chloroplast due to the fact that the PQ and NAD(P)H pools are connected by NDA2 (or indirectly, through Fd by the NDH in higher plants). Interestingly, the overreduction of the PSI donor- and acceptor sides in the ΔPTOX2 mutant at the onset of light was alleviated during the light period, as evidenced by an increased quantity of P700^+^ accumulated in the light and larger photo-oxidisable fraction of the P700 than in the WT at the end of the light interval.

Several hypotheses could account for the observed decrease in the PSII ETR during fluctuations: (i) PTOX2 plays a significant quinol oxidase role in the fluctuating light conditions; (ii) The overreduction of the downstream acceptors of PSII promotes photoinhibition which decreases PSII activity; (iii) parts or ensemble of the pool of PSI acceptors cannot be initially oxidised by CBB cycle, resulting in a block of electron transfer; (iv) supramolecular changes, mediated by the stromal redox state during the dark periods, prevent regular photosynthetic electron transfer.

Hypothesis (i) fails to account for the fact that maximal rate of PTOX is too low to produce significant differences in Q_A_ oxidation state; the hypothesis (ii) also can also be excluded in our conditions as the F_V_/F_M_ and ΦPSII decrease throughout the treatment is reversible in short timescales (see supplementary discussion for details). Hypothesis (iii) requires further assumptions to be taken into account. Considering a homogenous system in which all electron transfer chains are interconnected, then both NADPH and PQ pool are reduced upon illumination and subsequently re-oxidised by the PSI and CBB cycle, respectively, losing the “memory” of their respective initial states on the onset of illumination, unless a long-term effect on the CBB cycle kinetics was present in the chlororespiration mutant due to a transient overreduction. Nevertheless, there are several reports supporting the view that neither the thylakoid membranes, nor the stroma of the chloroplast of Chlamydomonas are homogenous (31), implying that pools of CBB cycle enzymes are connected exclusively with certain PSI centres. Moreover, although the exact concentration of PTOX is not known for Chlamydomonas, it is highly substoichiometric in plants with regards to PSII (32). Thus, even if the differences in the rates of PQ oxidation between algae and higher plants are a simple result of PTOX being more abundant in the former, it would still be less concentrated than the PSII. It can be excluded that microcompartmentation produces a constantly reduced fraction of the NADPH pool that lacks access to chlororespiration enzymes. Such a fraction would block entire units of electron transfer chains, lower the ΦPSII, and increase the reduction state of the P700. Contrary to this hypothesis, P700 oxidation measurements showed that the photo-oxidisable PSI fraction in the light at the end of the one-minute period of illumination was higher in the mutant. Our P700 measurements represent only an average state of the chloroplast in both strains. To explain the different fractions of photo-oxidisable P700, we favour a supramolecular reorganization of the photosynthetic apparatus in the PTOX mutant subjected to fluctuating light due to an overreduction of the chloroplast in darkness. One possibility for such regulation is the previously observed movement of cyt. *b*_6_*f* to the non-stacked regions of the thylakoid membrane linked to state transitions (33), although in our conditions no differences in the latter process were observed (Figs S2, S3B). However, this hypothesis could be *a priori* reconciled if the movement of cyt. *b*_6_*f* in reducing conditions is uncoupled from PSII/PSI antenna sharing (34).

Regardless of the reason why exactly the CEF operates at higher rates in the PTOX mutant, its comparison with the WT in fluctuating light proved well suited to study the regulation of CEF. Until noow, CEF regulation studies were hampered by anoxic conditions, which impeded LEF from water to CO_2_. Since ΔPTOX2 showed an increased CEF in fluctuating light suggests that the redox state of the chloroplast is a signal for the induction of CEF in physiologically relevant conditions (i.e. without DCMU).

Previous observations in anoxia (Nawrocki et al., *in preparation*)(35) showed increased CEF rates, contributing to a net ATP production which allow a more efficient consumption of NADPH by CBB cycle. However, here we provide evidence that in oxic conditions, chlororespiration is a critical regulator of the chloroplast redox poise in the dark, which results in its control over the redox state of both the PQ pool and the PSI stromal acceptors. It is however difficult to pinpoint the exact source of signalisation for CEF induction. Even though CEF is a PQ-reducing activity, it helps oxidising the PQ simply by increasing the rate at which NADPH is consumed (at least in conditions where the cyt. *b*_6_*f* is not limiting for electron flow). This implies that both of these pools could be a signal to increase CEF. Joliot & Joliot (36) also hypothesized that ATP/ADP ratio could be the sensor for CEF regulation.

In our experiments, CEF indeed was able to compete with LEF, suggesting that the two modes do not share the same pool of PQ or that a decrease in LEF enabled transitory, rapid CEF rates. Indeed, after establishment of a steady-state under fluctuating light (after 15 minutes), the ΔPTOX2 mutant displayed a ΦPSII which remained constant during the illumination while the fraction of photo-oxidisable PSI changed from ^~^0.3 to ^~^0.8. Because CEF is active during this period, as evidenced by the decrease in ΦPSII/ΦPSI ratio, and because the light we used for the treatment resulted in a major reduction of the PQ pool up to ^~^95%, it is likely that such an efficient CEF requires a heterogeneity in the plastoquinone distribution between those that are connected with PSII and those which are involved in CEF. Furthermore, a ^~^40% drop in the ΦPSII is also seen in the treatment with a five-fold increased light intensity (fig. S3), where over 98% of PQ is reduced. No standalone enzyme which mediates CEF as an Fd:PQ oxidoreductase has been found despite long-lasting efforts (Nawrocki et al., *in preparation*) (36-38). Thus, to account for its high rates (as discussed in Supplementary discussion), we would like to reiterate that CEF is mediated as initially proposed by Mitchell (39-41), via the haem c_i_ in cyt. *b*_6_*f* (42). In that respect, a plastoquinone released at the Q_o_ site could be used at the Q_i_ site for re-reduction without diffusing within the PQ pool exterior of the cyt. *b*_6_*f* dimer. Such a mechanism would allow for a fast CEF with rates of the same order of magnitude as LEF, as observed in higher plants and algae (Nawrocki et al., *in preparation*) (despite the PQ pool redox imbalance pointed out by Allen (43); but see discussion in (44)), and would account for the observation that PQ pool in our experimental conditions is reduced despite an oxidised PSI acceptor side and very rapid CEF (see also discussion in Nawrocki et al., *in preparation*).

As suggested earlier (45-47) by referring to the lateral heterogeneity along thylakoid membrane leading to a heterogeneity in the distribution of photosynthetic complexes, the diffusion of PQ is locally restricted (48, 49). Most of the PSI centres and roughly half of the cyt. *b*_6_*f* complexes are located in the non-appressed regions of thylakoids (50). This increases the probability that an oxidised quinone, despite having diffused out of the *b*_6_*f* dimer, is available for a hypothetical-FQR-mediated reduction. An increase in this spatial heterogeneity in fluctuating light could also account for the increased CEF rates.

If indeed the availability of oxidised quinones is not a limiting aspect for CEF, then the regulation should take place at the PSI acceptor side, and determine the probability with which the electrons return to quinones. Two major types of CEF structuration were proposed in the literature, either involving a differential binding of FNR to the cyt. *b*_6_*f* depending on the stromal redox state, or a closing in between PSI and cyt. *b*_6_*f*. (51, 52); as mentioned before, differences in the CBB cycle activity could also alter rates of cyclic in corresponding timescales and without a need of structural alternation in the thylakoids.

Our experiments in fluctuating light substantiate hypotheses that chlororespiration is a process regulating the redox state of the chloroplast stroma. This may explain the complex evolutionary history of PTOX enzymes that keep being duplicated, especially in microorganisms (12, 53) often exposed to fluctuations in light and carbon availability (54), or inhabiting dense populations with oxygen limitations (where PTOX could act as an oxygen sensor for ST, promoting State II in hypoxia to prepare the photosynthetic apparatus to the onset of light (35)). It is equally interesting to hypothesize about the rates of chlororespiration – why it is an order of magnitude lower in higher plants than in algae (16), contrary to cyanobacteria, which possess additional terminal oxidases (29). Plants have retained the complex I-like NDH as a PQ reductase, and it has been proposed that this enzyme is a Fd:PQ and not a NAD(P)H:PQ oxidoreductase (55) - the scarce amounts of its substrate in the dark result in a low reductive pressure on the pool and may explain the low abundance and/or rates of PTOX. However, despite those limitations, chlororespiration is surprisingly able to regulate the entire redox state of the stroma and PQ pool in darkness, which turns out to be crucial for photosynthesis during light fluctuations through changes in the LEF to CEF partitioning.

## Materials and methods

### Strain and growth conditions

For all the experiments, a descendant strain of C. reinhardtii cc137 was used. Prior to the project, the ΔPTOX2 mutant (20) was back-crossed with the reference strain to keep them genetically close. The strains were grown in 25° C in tris-acetate-phosphate (TAP, mixotrophic) or minimum (autotrophic) media in continuous continuous ^~^10 μE.m^-2^.s^-1^ light, unless stated differently.

### Growth measurements

The growth of the WT and ΔPTOX2 mutant simultaneously grown in auto- and mixotrophic conditions in a culture chamber (Infors HT, Switzerland) with a custom controlled, white LED illumination system (BeamBio, France) for at least 72 hours was measured in three repetitions using a cell counter (Beckman Coulter, USA). The growth rate was fitted with an exponential (OriginLab software) yielding the rate in percent of cell multiplication per hour. Because the scatter of absolute growth rates between biological replicates was larger than the relative differences of the growth rate between the WT and the mutant in a single experiment, while the relative decrease of growth in the mutant was virtually constant, the growth of the latter is expressed as relative to the WT.

### Functional measurements

Prior to experiments, apart from ΦPSII measurements directly from the fluctuating light cultures, the cells were harvested by centrifugation (3000 rpm, 5 min) resuspended in minimum medium supplied with 10% Ficoll to avoid excessive sedimentation. They were then adapted for 1h to low light intensity (^~^10 μE.m^-2^.s^-1^) in order to set both of the strains in state I.

The electron transfer rate from PSII (ETR PSII) was calculated as the light intensity × ΦPSII × σPSII or I × ΦPSII after 30 s adaptation to a given light, where σPSII is the absorption cross-section of PSII. The intensity I (light intensity × σPSII) is expressed as e^-^.s^-1^.PSII^-1^ thanks to measurements of the area above fluorescence induction curve in the presence of DCMU where only the single electron (P_680_ to Q_A_) transfer occurs (56).

Other fluorescence kinetics and P_700_ redox measurements were performed in the JTS-10 spectrophotometer (BioLogic, France) as described before (5), except that they were conducted in an open cuvette setup, with regular air-bubbling of the sample to avoid anoxia and sedimentation of the cells during the 45 min treatments.

The PSI parameters were calculated as follows: ΦPSI = (maximal photo-oxidisable P_700_)-(P_700_ in the light); photo-oxidisable P_700_ = (maximal photo-oxidisable P_700_)-( P_700_ in darkness); and P_700_^+^ accumulated in the light = (P_700_ in the light)-( P_700_ in darkness). They were all reported as a fraction of total P_700_^+^: total P_700_^+^ = (maximal photo-oxidisable P_700_ in the presence of DCMU)- (P_700_ in darkness). The maximal values ox photooxidable P_700_ were obtained using a saturating pulse of ^~^3000 μE.m^-2^.s^-1^ in the presence of DCMU.

Fluorescence emission spectra at 77K were recorded as described previously (5).

### Biochemistry

To investigate whether a significant proportion of PTOX2 becomes stroma-soluble in the darkness as proposed before we have separated the soluble- and membrane fractions of cells grown to early log phase. The cells were harvested, concentrated and exposed to 1 hour continuous moderate light (50 μE.m^-2^.s^-1^) or oxic darkness treatment. They were then broken using French’s press and spun down to remove unbroken cells. The supernatant was ultracentrifuged for 20 minutes and the resulting supernatant was used as soluble fraction. The thylakoid membranes were then isolated as previously described (34), separated by SDS-PAGE and Western blotted. The blotting was repeated until a correct adjustment of respective protein markers in the fractions was achieved.

In order to measure the accumulation of PTOX2 in different conditions, the cells were grown to early log phase in low light (^~^10 μE.m^-2^.s^-1^) and shifted to high light (const. 300 μE.m^-2^.s^-1^) of fluctuating light (60 s / 60 s, 0/300 μE.m^-2^.s^-1^) for 5 or 24 hours. They were then harvested and whole cell protein extracts were loaded on chlorophyll basis and separated using a 12% SDS-PAGE.

## Acknowledgements

We thank Stefano Santabarbara for experiment suggestions. Sandrine Bujaldon is acknowledged for a cross between the WT and the ΔPTOX2 mutant. W.J.N. had support of the French Ministry of Education. The project was supported by the French state (Labex DYNAMO ANR-11-LABX-0011-01).

## Supplementary figures

**Figure S1.**
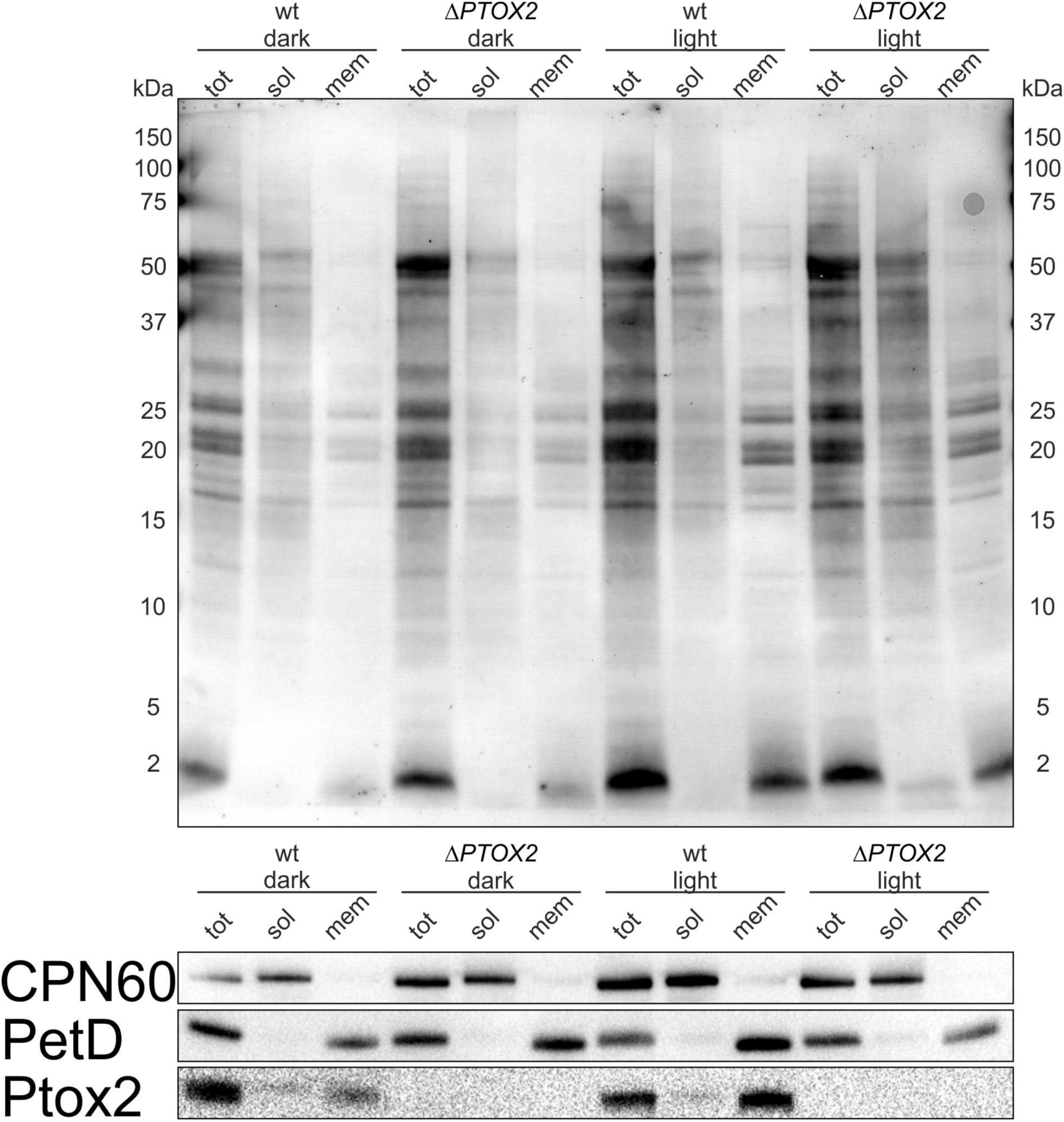
Fractionation of broken Chlamydomonas cells. The strains from continuous light and darkness were broken in a French press (total sample), and membranes after ultracentrifugation were floated on step sucrose gradient (membrane fraction). Soluble fraction is the supernatant of ultracentrifuged total sample. CPN60 is a soluble, major chloroplast chaperone.

**Figure S2.**
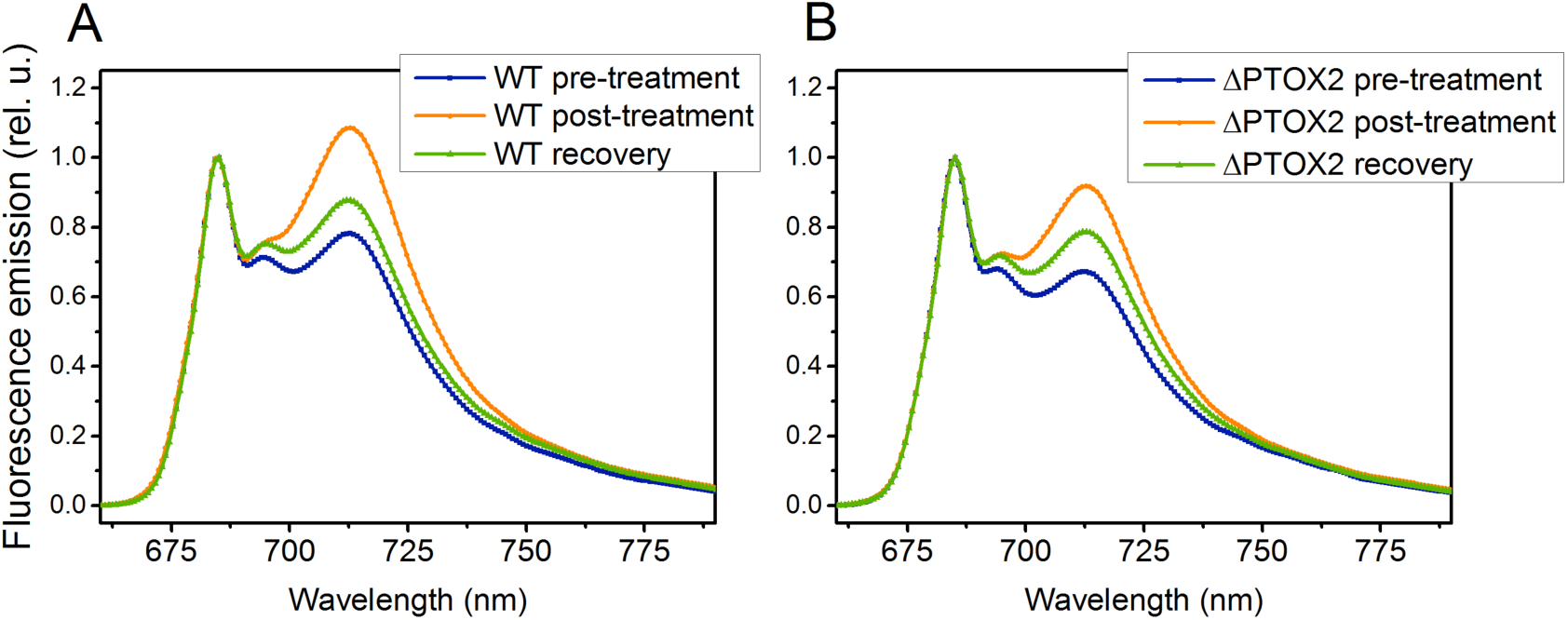
77K fluorescence emission spectra of the WT (A) and the PTOX2 (B) mutant grown in mixotrphic conditions with continuous illumination. The spectra were measured before (“Pre-“) and immediately after (“Post-“) the 30 min. fluctuating light treatment. “Recovery“corresponds to posttreatment cells exposed to 15 mins of continuous low light for recovery.

**Figure S3.**
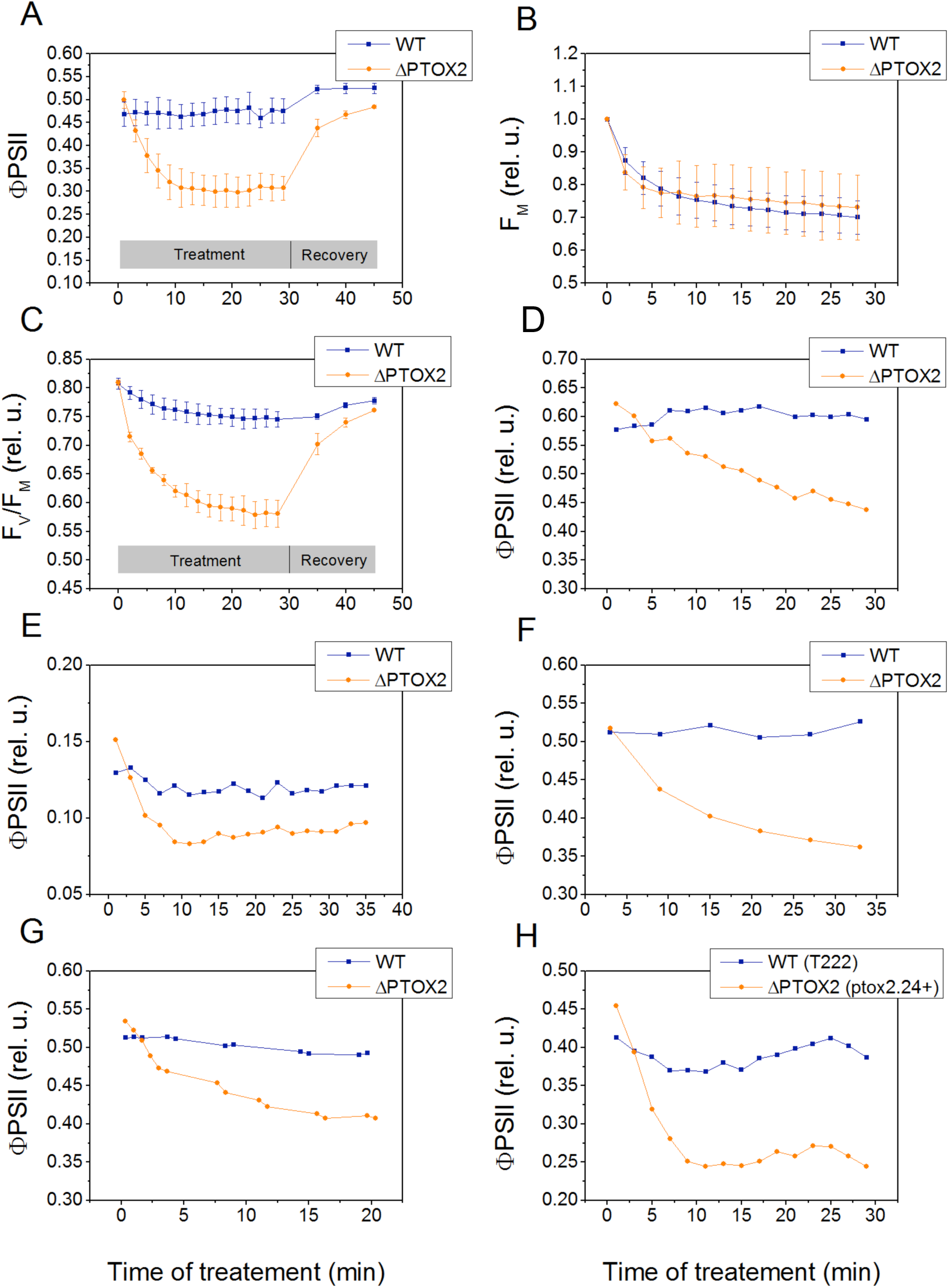
Room temperature fluorescence parameters of various fluctuating light treatments in the WT and the ΔPTOX2 mutant. A, ΦPSII during 60/60 s, 0/340 mE/m^2^/s treatment and low light recovery. B, F_Max_ during 60/60 s, 0/340 mE/m^2^/s treatment. C, F_V_/F_M_ during 60/60 s, 0/340 mE/m^2^/s treatment. D, ΦPSII during 60/60 s, 0/125 mE/m^2^/s treatment. E, ΦPSII during 60/60 s, 0/1500 mE/m^2^/s treatment. F, ΦPSII during 180/180 s, 0/340 mE/m^2^/s treatment. G, ΦPSII during 20/20 s, 0/340 μE/m^2^/s treatment. H, ΦPSII during 60/60 s, 0/340 mE/m^2^/s treatment in cells with different genetic background.

**Figure S4.**
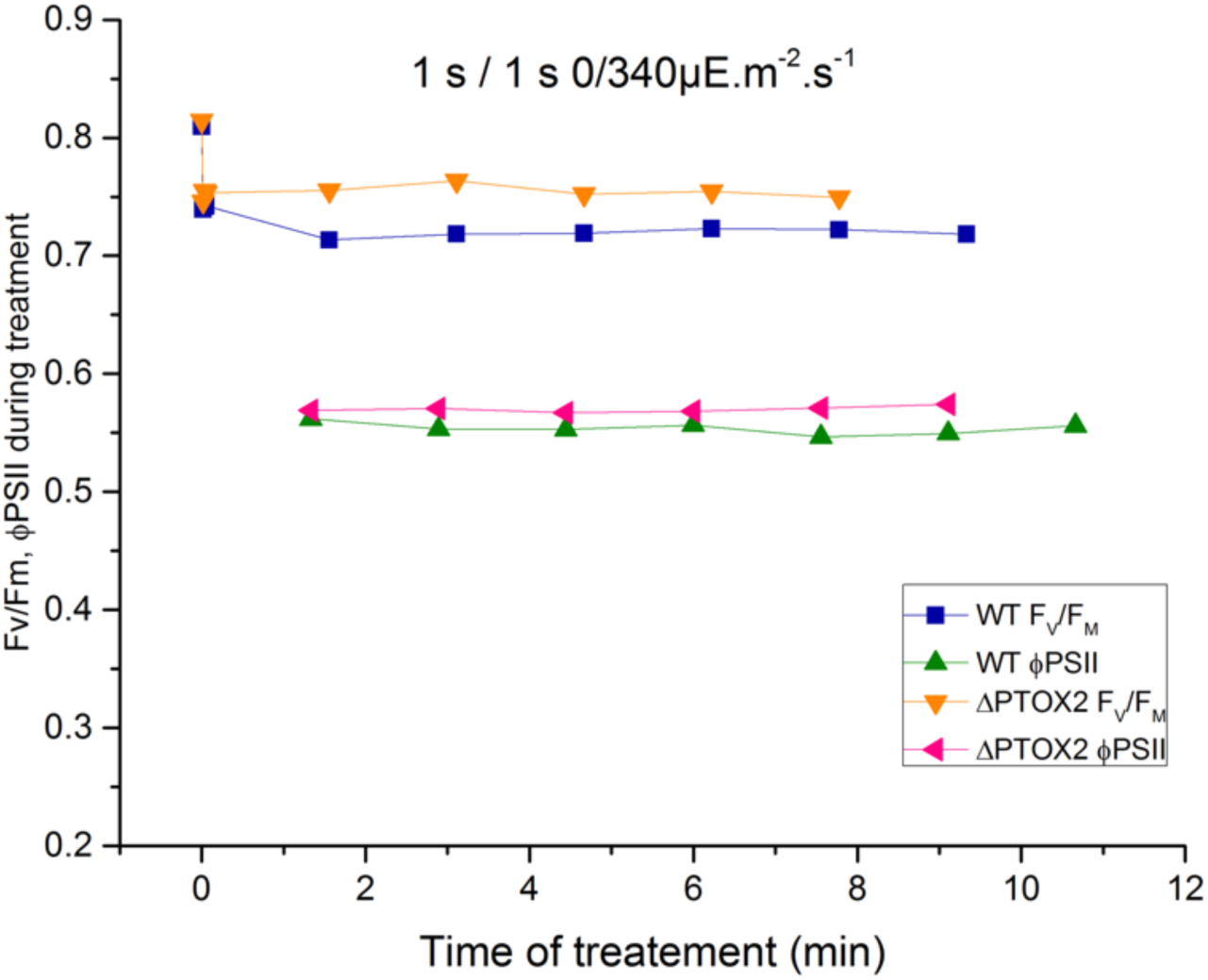
Room temperature ΦPSII and F_V_/F_M_ fluorescence parameters of 1/1 s fluctuating light treatment in the WT and the ΔPTOX2 mutant.

**Figure S5.**
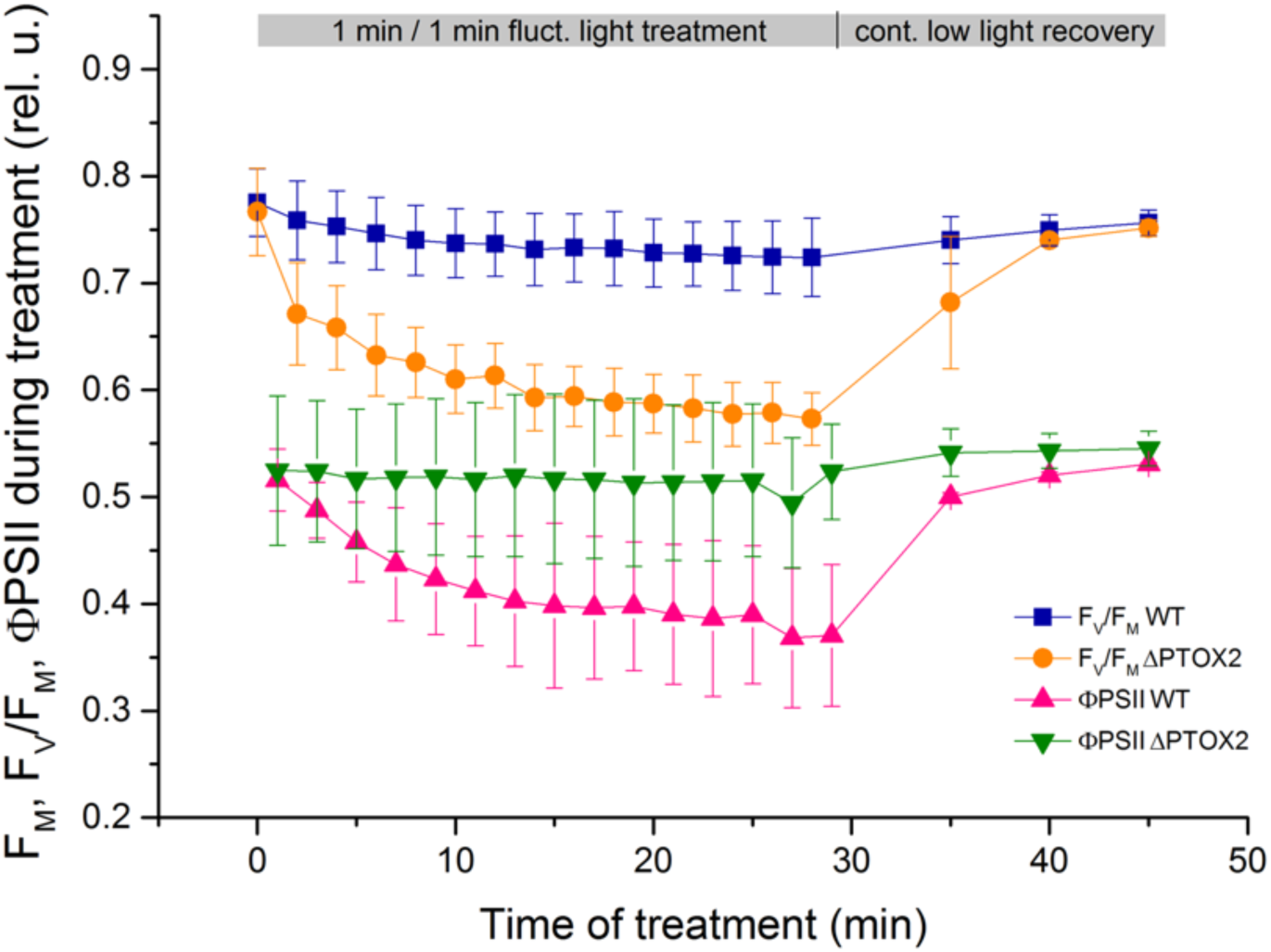
Room temperature ΦPSII and F_V_/F_M_ fluorescence parameters of 60/60 s fluctuating light treatment in the WT and the ΔPTOX2 mutant grown in autotrophic conditions.

**Figure S6.**
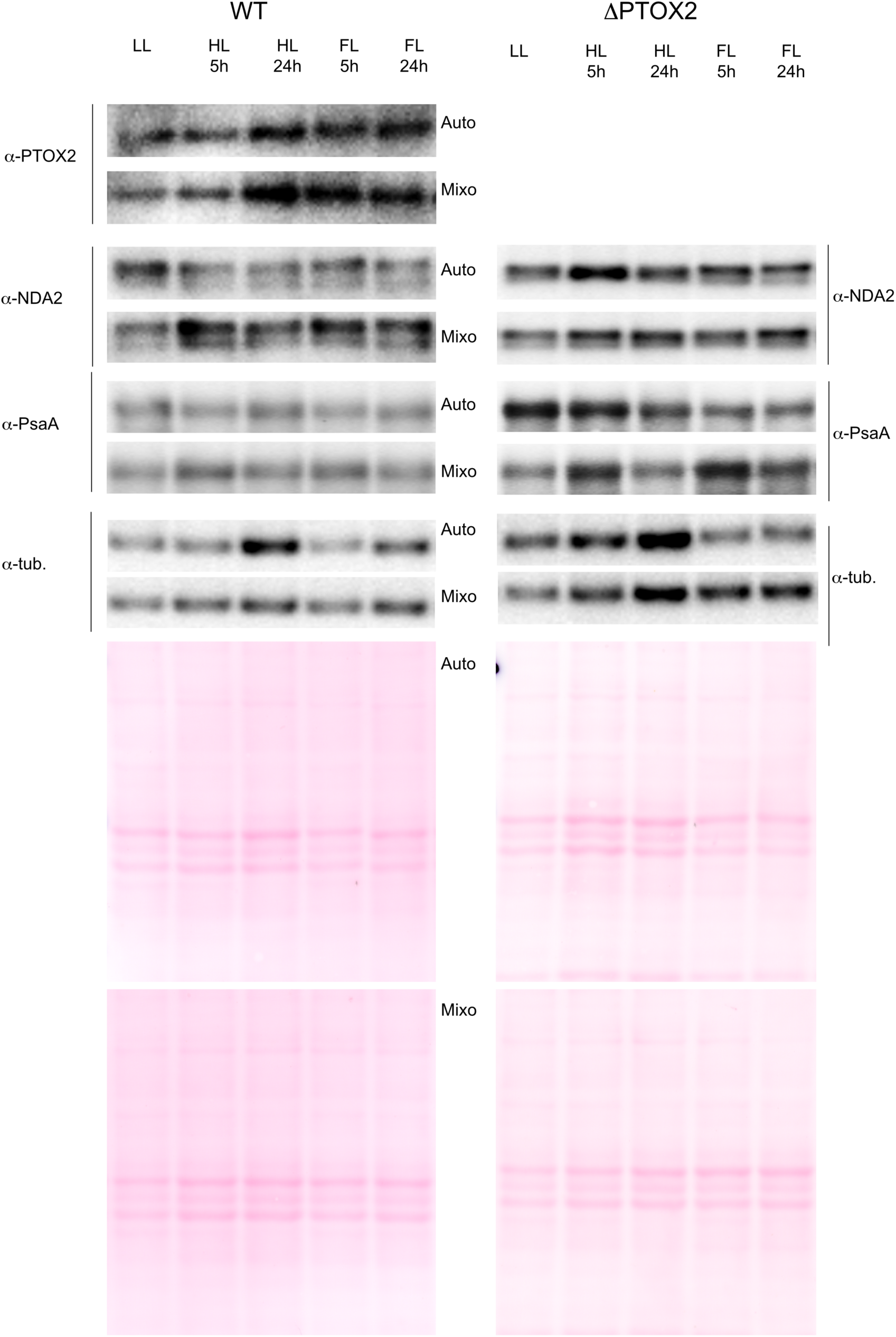
Accumulation of proteins in the WT and the ΔPTOX2 mutants in cont. low light (LL) and after 5- and 24 h exposure to cont. high light (HL) or fluctuating light (FL) in both auto- and mixotrophic conditions.

Supplementary methods
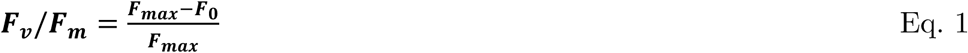

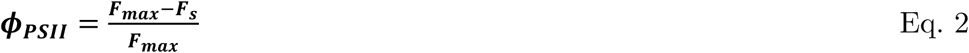

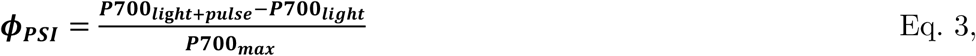

where ***P700_max_*** is the value upon saturating pulse in a DCMU-treated sample.

Note that the ***P700_light+pulse_*** value is lower due to acceptor side limitation of

P700.
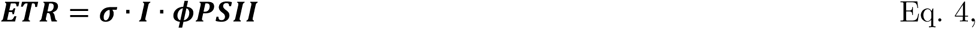

where ***σ*** is the cross-section of PSII and ***I*** the light intensity.

According to Stern-Volmer relationship, quencher concentration influences the rate of fluorescence in a following manner:

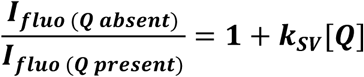

where ***I_fluo(Q absent)_*** is the fluorescence intensity without a quencher, ***I_fluo(Q present)_*** with the quencher, ***K_sv_*** is the quenching coefficient and **[*Q*]** the concentration of a quencher.

A transformation of this equation gives:

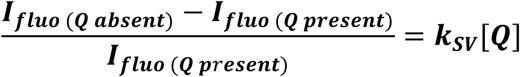

One can substitute ***I_fluo(Q absent)_*** to ***F_max_*** (the maximal level of fluorescence when the Q_A_ is in its reduced state after a saturating pulse), ***I_fluo(Q present)_*** to ***F***, and the **[*Q*]** to **[*Q_a(ox)_***], which yields:

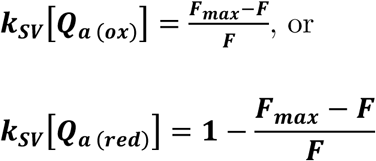

Supplementary discussion:

Because the ΦPSII and ΦPSI parameters provide photochemical yields rather than electron transfer rates (ETR) by respective photosystems, we have tried to estimate the latter. ETR is a product of photochemical yield, absorption cross-section of the Photosystem and the light intensity. As the latter did not change during our experiment, it is important to investigate whether antenna distribution, known to be affected in the ΔPTOX2 mutant, can buffer or increase ETR values with regards to photochemical yields.

We have taken care to set both of the strains to similar state before the treatment. This is evidenced by high and identical values of the first recorded F_V_/F_M_ parameter (fig. S3 and S5) and very similar, state-I-like fluorescence emission spectra before the treatment (fig. S2). This way, we eliminate the problem occurring by dark-adapting the PTOX2 mutant prior to the experiments, which would set it in state II, and we introduce an oxidized initial state of the cells (F_V_/F_M_ higher in both strains than in dark-adapted WT).

There are several indications that the changes of antenna, initially in the same state, are changing in a similar way. The first indication is the F_M_ parameter, and its decrease which is identical and limited in both cases (fig. S3). Changes of F_M_ can indicate two things, state transitions or NPQ. Let us consider two borderline scenarios, where the changes in F_M_ are i) entirely explained by ST, and ii) completely due to NPQ, and how the two would influence the ETR.

In the first case scenario, a decrease of F_M_ indicates transition from state I (or a condition close to state I) to state II, so a transfer of LHCII antenna from PSII to PSI (5). In such case, the observed ^~^20% F_M_ decrease during the treatment corresponds to a decrease of PSII-, and an increase of PSI antenna size of 20%. Because the respective photochemical yields change in the same direction, i.e. ΦPSII decreases and ΦPSI increases, the resulting PSII ETR would decrease much more than the ΦPSII indicates (and PSI ETR would correspondingly be underestimated by investigating only ΦPSI).

In the second case, ii), NPQ would be the sole reason for the F_M_ decrease. It is possible that some slowly-relaxing NPQ occurs during the treatment, and it is not distinguishable from ST in the 77K emission spectra. The F_M_ decrease of 20%, however, having calculated it for the concentration of the quenching species with the Stern-Volmer relationship, would be a result of a functional inactivation of only 4% of PSII centers (and would not have an influence on PSI). Therefore, in the case ii), only a minor correction would need to be applied in order to translate the ΦPSII to a relative ETR, and the PSII ETR would still decrease. We have therefore proven that the both the initial state and antenna changes are identical for both strains, and that the latter would not compensate for differences we observe between the strains, but rather enhance them toward an even bigger discrepancy between the PSI and PSII fluxes in the mutant in fluctuating light.

We have also estimated the combined ETRs of both photosystems at the beginning- and at the end of the fluctuating light treatment. In the case ii), if solely NPQ explains F_M_ and cold emission spectra shifts during the treatment, the relative F values can be approximated as relative ETRs. If, however, ST are occurring during the treatment, the yields need to be corrected to yield relative ETRs:

**Supplementary Table 1.** Influence of the treatment on the photochemical rates (expressed in e^-^.s^-1^.2PS^-1^) in WT and DPTOX2.

| | WT | ΔPTOX2 |
| --- | --- | --- |
| Beginning of the treatment | 210 | 204 |
| End of the treatment | 224 | 176 |

As shown before, a transition from a quasi-perfect state I (PSI:PSII antenna size ratio of 1:1) to state II occurs during the treatment. As the F_M_ changes are complementary to the increase of PSI antenna size, we roughly estimate that the final ratio of functional size of the photosystems is therefore 1.2:0.8, (PSI antenna 50% bigger than PSII). Hence, the final ΦPSII for both strains should be ideally multiplied by 0.8, and the final ΦPSI by 1.2 to yield the fraction of contribution to the ETR. Roughly, this corresponds for the WT to ΦPSII of 0.38, ΦPSI of 0.57; ΦPSII of 0.23 and ΦPSI of 0.69 in the ΔPTOX2. These relative contributions to the photochemistry may be then used to precisely describe the ETR of each photosystem, including cyclic electron flow which contributes to PSI, but not PSII ETR: knowing that the photochemical rate in the WT is 220e/s/2PS and 170e/s/2PS in the ΔPTOX2 at the end of the treatment: PSII ETR is 88e/s, and PSI 132e/s (CEF of 44e/s) in WT, and PSII ETR is 43e/s and PSI ETR is 128e/s (CEF of 85e/s).

